# Disentangling the causes for faster-X evolution in aphids

**DOI:** 10.1101/125310

**Authors:** J Jaquiéry, J Peccoud, T Ouisse, F Legeai, N Prunier-Leterme, A Gouin, P Nouhaud, JA Brisson, R Bickel, S Purandare, J Poulain, C Battail, C Lemaitre, L Mieuzet, G Le Trionnaire, JC Simon, C Rispe

## Abstract

Faster evolution of X chromosomes has been documented in several species and results from the increased efficiency of selection on recessive alleles in hemizygous males and/or from increased drift due to the smaller effective population size of X chromosomes. Aphids are excellent models for evaluating the importance of selection in faster-X evolution, because their peculiar life-cycle and unusual inheritance of sex-chromosomes lead to equal effective population sizes for X and autosomes. Because we lack a high-density genetic map for the pea aphid whose complete genome has been sequenced, we assigned its entire genome to the X and autosomes based on ratios of sequencing depth in males and females. Unexpectedly, we found frequent scaffold misassembly, but we could unambiguously locate 13,726 genes on the X and 19,263 on autosomes. We found higher non-synonymous to synonymous substitutions ratios *(dN/dS*) for X-linked than for autosomal genes. Our analyses of substitution rates together with polymorphism and expression data showed that relaxed selection is likely to contribute predominantly to faster-X as a large fraction of X-linked genes are expressed at low rates and thus escape selection. Yet, a minor role for positive selection is also suggested by the difference between substitution rates for X and autosomes for male-biased genes (but not for asexual female-biased genes) and by lower Tajima’s D for X-linked than for autosomal genes with highly male-biased expression patterns. This study highlights the relevance of organisms displaying alternative inheritance of chromosomes to the understanding of forces shaping genome evolution.

## Introduction

Sex chromosomes are major players in evolution. Besides their role in sex determination, sex chromosomes contribute to genomic conflicts (Rice 1984; Meiklejohn and Tao 2010; Soh et al. 2014), genetic incompatibilities and reproductive isolation (Coyne and Orr 2004; Saether et al. 2007; Kitano et al. 2009; Johnson and Lachance 2012). A pair of sex-determining chromosomes typically evolves toward reduced recombination rates (crossing overs), which eventually causes one of the sex chromosomes to gradually lose most of the chromosomal regions (loci) present in the alternate one (Charlesworth et al. 2005). These loci will thus be found in single copy in the sex that carries the degenerate, smaller sex chromosome. When the heterogametic sex is the male, sex chromosomes are denoted X and Y (e.g., in mammals), whereas when it is the female, sex chromosomes are noted W and Z (e.g., in birds). Alleles of loci present only on the X (W) are more exposed to selection in individuals of the heterogametic sex, facilitating the fixation of beneficial mutations and the purging of deleterious ones (Charlesworth et al. 1987). On the other hand, because males (XY) bear and transmit a single X chromosome, effective population size is smaller for the X compared to autosomes (Wright 1931; Caballero 1994; Caballero 1995). This increases the rate of fixation of slightly deleterious mutations on the X by genetic drift (Kimura 1983) (the same principles apply to ZW systems, so we ignore these in the following). Consequently, X-linked genes may evolve faster than autosomes (“faster-X” evolution) due to higher positive selection (rate of fixation of beneficial mutations) and/or higher genetic drift (rate of fixation of slightly deleterious mutations) (Vicoso et al. 2009, Mank et al. 2010). Faster evolution of X-linked proteins is supported by observations in a large panel of species (e.g., *Drosophila*, nematodes, mammals, birds, see Meisel and Connallon 2013 for a review). In some, a preponderant effect of drift was demonstrated (Mank et al. 2010; Avila et al. 2014), while positive selection would play a predominant role in other species (Baines et al. 2008; Hvilsom et al. 2012; Langley et al. 2012; Mackay et al. 2012; Kousathanas et al. 2014; Sackton et al. 2014; Avila et al. 2015).

In this context, organisms with atypical inheritance of sex chromosome can greatly facilitate inferences about the processes contributing to the evolution of sex chromosomes (Bachtrog et al. 2011). Among those, aphids, which present X0 males and XX females, have equal effective population sizes for X chromosomes and autosomes (Jaquiéry et al. 2012a). This eliminates one possible confounding factor of faster-X evolution, the smaller effective population size of the X, helping to properly test its causes. Aphids reproduce by cyclical parthenogenesis, such that males and sexual females constitute only a short part of their life-cycle, which is dominated by apomictic parthenogenetic (clonal) XX females (figure 1).

**Fig. 1.**
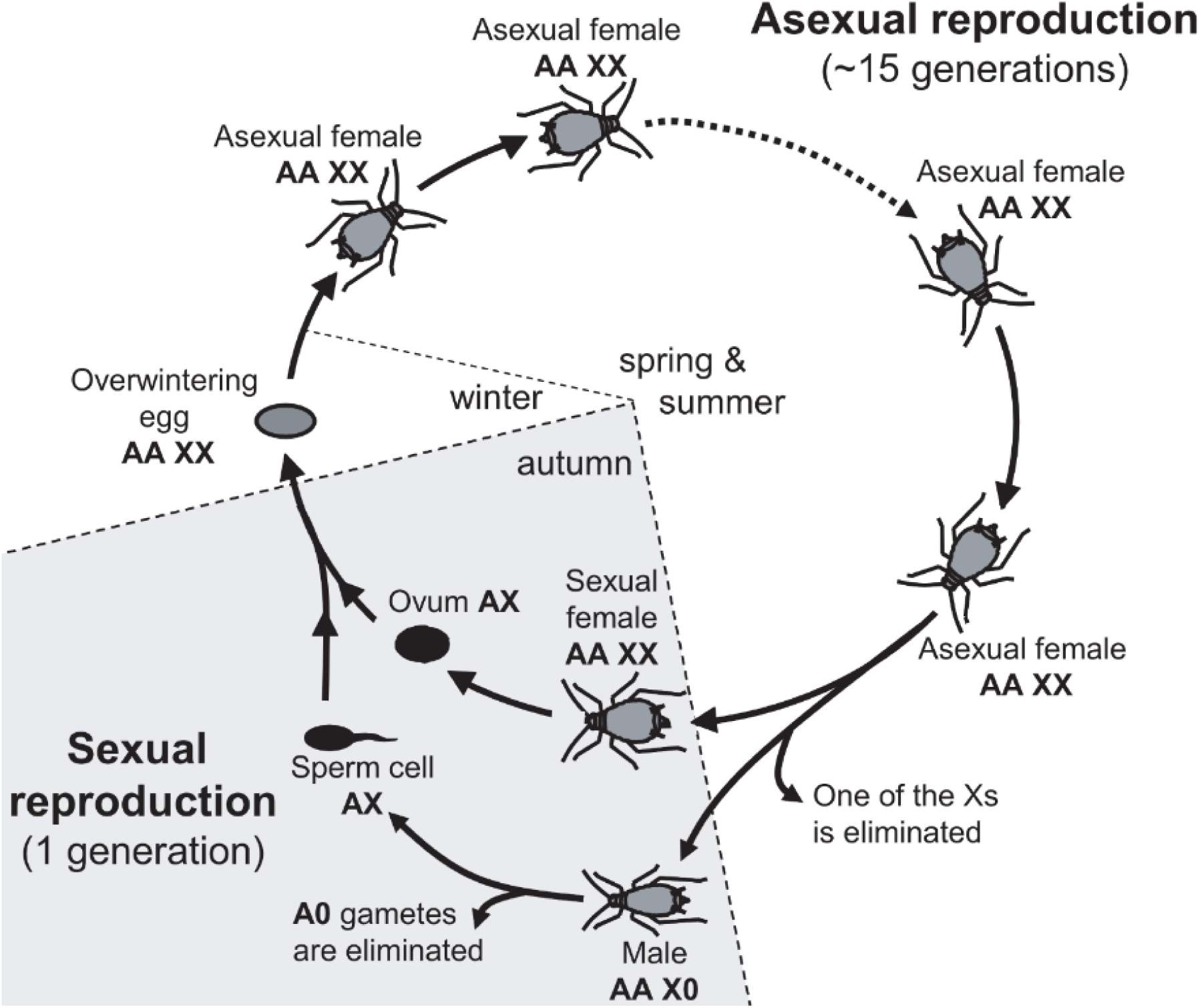
Life-cycle of the pea aphid and ploidy levels for autosomes (A) and the sex-chromosome (X) (adapted from Jaquiéry et al. 2013).

Males are produced asexually via the elimination of one X from the germ line (Wilson et al. 1997; Caillaud et al. 2002). As a result, X-linked recessive alleles are exposed to selection in male aphids, just like in other X0 or XY males. However, because all sexually-produced aphid eggs are XX females, all the progeny inherit their X from males and sexual females in equal proportions. This difference from other heterogametic systems, where progeny present even sex ratios, has deep consequences for the evolutionary trajectory of the aphid X chromosome (Jaquiéry et al. 2012a). Furthermore, in contrast to standard systems, variance in reproductive success between sexes, population expansion, bottlenecks and sex-biased dispersal should not differentially affect sex chromosomes and autosomes in aphids (Jaquiéry et al. 2012a). Mutation and recombination rates are also expected to be equal across chromosomes because of their similar mode of inheritance and the complete absence of crossing overs in males (Jaquiéry et al. 2012a). These similarities between X chromosomes and autosomes make aphids exceptionally useful to pinpoint the causes of faster-X evolution, since the factors mentioned above need not be accounted for. Still, a notable difference between X and autosome in aphids is the theoretical propensity of the X to accumulate sexually antagonistic mutations beneficial for males and detrimental to asexual females, which also is the consequence of cyclical parthenogenesis combined with the inheritance of the X (Jaquiéry et al. 2013).

Empirical analyses on a small subset of genes of the pea aphid (*Acyrthophon pisum*) showed that X-linked genes evolve faster than autosomal genes (Jaquiéry et al. 2012a) and that genes expressed predominantly in males (hereafter “male-biased” genes) predominantly locate on the X (Jaquiéry et al. 2013). Subsequent genome-wide analyses did not however support faster-X evolution (Purandare et al. 2014), and mitigated the degree of enrichment of male-biased genes on the X (Purandare et al. 2014; Pal and Vicoso 2015). These discrepancies however likely stem from the fact that these two studies did not assign individual genes to chromosome types, but entire scaffolds, which present assembly errors (as shown by Bickel et al. 2013 and suggested by Jaquiéry et al. 2013). Misassignment of genes to chromosomes would indeed artificially decrease the contrast between X-linked and autosomal genes.

Here, we aimed to go past these shortcomings in order to fully disentangle the causes for faster-X evolution in aphids. For this, we first attributed genes to the X or to autosomes at the scale of the entire genome in the pea aphid. On a large set of genes, we combined estimates of substitution rates at the interspecific level with polymorphism data in pea aphid populations and gene expression levels in the various genders and morphs. This allowed assessing how relaxed selection (genetic drift) and adaptation contribute to the faster evolution of the X chromosome in this system.

## Material and Methods

### Assignment of scaffold regions to the X and autosomes

#### Full-genome sequencing of females and males

An asexual aphid mother has the same diploid autosomal genome as her sons, but has two X chromosomes instead of just one (figure 1). We took advantage of this XX/X0 system to assign the pea aphid genome sequence (Acyr 2.0, Genbank accession GCA_000142985.2, IAGC 2010) to the X or to the autosomes by comparing sequencing depth along assembled scaffolds between mapped reads from females and from males of the same parthenogenetic lineage (clone). DNA from five asexual females and five winged and five wingless males of clone P123 (Simon et al. 2011) was extracted with the Qiagen DNeasy Blood and Tissue kit, following the manufacturer’s protocol. Wing male polymorphism in this clone was used to determine the X copy that each male carried, based on the knowledge that the locus that controls this trait is X-linked (Caillaud et al. 2002) and is heterozygous in clone P123 (Frantz et al. 2010). Each individual was genotyped at seven polymorphic microsatellite markers (Peccoud et al. 2008) to confirm its identity. One of those markers, which is known to be X-linked (Caillaud et al. 2002), allowed us to confirm the nature of the X copy inherited by each male. Three DNA extracts were obtained: one from five females and one per genotype of males, using five individuals in each. These samples were sequenced on the Illumina Hiseq 2000 platform yielding 100 bp pair-end reads at ∼43X coverage for the females sample and 25 to 30X coverage for each male type. Reads from each sample were mapped on scaffolds of the pea aphid genome assembly (Acyr 2.0) and on genome sequences of the bacterial symbionts of this pea aphid clone using the method described in Gouin et al. (2015) using Bowtie 2 (Langmead and Salzberg 2012) with proper insert sizes and parameters set as default. Depth of coverage at each nucleotide position of the reference genome was recorded and single nucleotide polymorphisms (SNPs) were identified using GATK’s Haplotype Caller (McKenna et al. 2010; DePristo et al. 2011). The raw sequence data has been deposited in the SRA division of Genbank (project accession: ERP022905 and PRJNA385573).

#### Comparison of sequencing depth between males and females

The following analysis was performed in R (R Development Core Team 2015). We analyzed genome positions covered by 20 to 70 reads in the asexual female sample, a range chosen to eliminate regions with low-coverage and regions with suspiciously high coverage (potentially duplicated or repeat-rich regions). Since overall coverage was slightly higher for one of the male types (∼30X) than for the other (∼25X), we normalized the depth of coverage data of the second male type (multiplying coverage estimates by a 30/25 ratio). We then averaged depth of coverage at each base position over male types. The ratio of median coverage depth of males to median coverage depth of the female sample was calculated on 10-kilobases (kb) scaffold windows sliding by 2-kb steps. A single window was used for scaffolds shorter than 10 kb. We expect the ratio of median coverage depth to be twice larger for autosomal regions than for the X chromosome. Accordingly, this ratio had a clearly bimodal distribution (supplementary figure S1), with modes at 0.34 and 0.66. We assigned a 10-kb window to the X if its ratio ranged between 0.2 and 0.445, to autosomes if it ranged between 0.53 and 1, whereas the region was tagged as “ambiguous” if it ranged between 0.445 and 0.53. Windows assigned to the same chromosome type and which were separated by less than four consecutive “ambiguous” windows were aggregated into a scaffold region we call a “block”. A whole block, including its “ambiguous” windows, was assigned to the corresponding chromosome type.

#### Comparison of male and female genotypes

Inheritance of single nucleotide polymorphisms (SNPs) also informs on the type of chromosome carrying a scaffold block. In fact, SNPs that are heterozygous in females but are also heterozygous in males are necessarily located on autosomes. Conversely, SNPs which are heterozygous in females but homozygous in males must be on the X. This SNP-based approach is however expected to be less powerful than the depth of coverage-based method for genomic regions with low heterozygosity. Thus, we only used SNP data to validate X/A assignments based on depth of coverage ratio (supplementary figure S2). A position was determined as heterozygous if the rarest allele was represented in at least 25% of reads, otherwise it was considered as homozygous. Assignment of SNPs to chromosome types was performed according to the genotypes of males, as described above. SNPs showing inconsistent genotypes (e.g., females and males of one type are both heterozygous while males of the other type are homozygous) were not assigned. SNP-based assignments were then visually compared to assignments based on depth of coverage (supplementary figure S2).

### Assignment of predicted genes to the X and autosomes

We used the 36,990 genes that constitute the gene prediction v2.1 for the Acyr 2.0 genome assembly available at http://bipaa.genouest.org/is/aphidbase/. Each of these genes was determined as X-linked or autosomal if the full length of its coding sequence (CDS) was comprised in a single scaffold block or was spread over several scaffold blocks assigned to the same type (either X, or A). Genes that could not be unambiguously assigned (mainly because they located on “ambiguous” blocks) were removed from further analyses. We also excluded 589 predicted genes that corresponded to rRNA (non-coding DNA).

### Sex-biased gene expression

We used the eight RNAseq libraries from Jaquiéry et al. (2013) to characterize gene expression patterns between morphs. Briefly, these eight libraries correspond to whole insects, with three male libraries, three parthenogenetic female libraries and two sexual female libraries – different libraries in each morph representing biological replicates – using adults of a single clone of *A. pisum* (clone LSR1). Details regarding aphid rearing, library preparation and sequencing are provided in Jaquiéry et al. (2013). Libraries were mapped on Acyr 2.0 as described previously. The number of reads covering each CDS was then counted. Read counts were normalized with the R package DESeq with default parameters (Anders and Huber 2010). For each gene, the effect of the morph (a three-level factor comprising male, sexual female and asexual female) on expression was tested with a GLM (R package MASS, Venables et al. 2002) with a quasi-poisson distribution of residuals, considering the different libraries for each morph as replicates. *P*-values were corrected for multiple testing using the Benjamini-Hochberg method implemented in R. Genes differentially expressed between morphs (*p*<0.05 after adjusting for multiple testing) were then categorized according to their pattern of expression in the different morphs as described in table 1.

**Table 1.** Number of X-linked and autosomal genes and frequency of X-linkage for classes of genes with contrasted patterns of expression between morphs.

| Category of genes | number of X-linked genes | number of autosomal genes | Frequency of X-linkage | *p* value <sup>6</sup> |
| --- | --- | --- | --- | --- |
| All<sup>1</sup> | 13726 | 19263 | 0.42 | 10<sup>-16</sup> |
| Low expression<sup>2</sup> (less than 10 reads/kb) | 10995 | 8136 | 0.57 | 10<sup>-16</sup> |
| Expressed<sup>3</sup> (at least 10 reads/kb) | 2771 | 11127 | 0.20 | 0.0001 |
| Unbiased<sup>4</sup> (>10 reads/kb & padj>0.1) | 697 | 3355 | 0.17 | na |
| 2-fold male-biased<sup>5</sup> | 1546 | 2245 | 0.41 | 10<sup>-16</sup> |
| 5-fold male-biased<sup>5</sup> | 962 | 948 | 0.50 | 10<sup>-16</sup> |
| 2-fold sexual female-biased<sup>5</sup> | 448 | 1369 | 0.25 | 10<sup>-10</sup> |
| 5-fold sexual female-biased<sup>5</sup> | 148 | 407 | 0.27 | 10<sup>-7</sup> |
| 2-fold asexual female-biased<sup>5</sup> | 244 | 1023 | 0.19 | 0.10 |
| 2-fold asexual female-biased<sup>5</sup> | 93 | 423 | 0.18 | 0.68 |

<sup>1</sup>all predicted genes that were assigned to the X or autosomes are included
<sup>2</sup>genes with on average <10 reads/kb of exon (average over the 3 morphs)
<sup>3</sup>genes with on average ≥10 reads/kb of exon (average over the 3 morphs)
<sup>4</sup>genes with on average ≥10 reads/kb of exon (average over the 3 morphs) and with an adjusted p-value ≥ 0.1 when tested for morph-biased expression.
<sup>5</sup>A gene was included in the morph-biased category (either male-, female- or asexual-biased) if the adjusted p-value for a morph effect was <0.05 and if it was at least x-fold (2 or 5) more expressed in one of the morph compared to the two other morphs.
<sup>6</sup>Deviation from expectation (given by the “unbiased” category) was evaluated with a test of proportion.

### Evolutionary rates

To assess substitution rates in X-linked and autosomal genes, sequences from another aphid species were necessary. *Acyrthosiphon svalbardicum* was chosen, ensuring sufficient genetic divergence, and reducing the risk of mutational saturation and of chromosomal rearrangements between the two species. Asexual females of *A. svalbardicum* were collected in Svalbard in 2009, and were then reared in the lab under 10:14 light:dark and 15°C on *Dryas octopetala*. Ten females were then frozen into liquid nitrogen and kept for subsequent RNA extraction performed using the RNeasy plant mini kit (Qiagen) according to manufacturer’s instructions. Two separate RNA extractions of 5 adults were performed. RNA quality was checked on Bioanalyzer (Agilent) and quantified on Nanodrop (Thermo Scientific). One sample made of a pool of 2 μg of the two independent RNA extractions was subsequently sent to GATC Company for RNA paired-end sequencing. The raw sequence data has been deposited in the SRA division of Genbank (project accession: PRJNA385897).

A *de novo* transcriptome assembly for *A. svalbardicum* was obtained following the methods of Rispe et al. (2016). Low quality parts of the reads were trimmed from the right ends with prinseq-lite (http://prinseq.sourceforge.net/) when the mean of phred score in a 20-bp window was below 20. Reads longer than 20 bp after trimming were re-organized by pairs (orphans were suppressed) and assembled with Trinity (Grabherr et al. 2011) using default parameters. Coding regions were predicted using FrameDP (Gouzy et al. 2009). Reciprocal BLASTN (Altschul et al. 1990) searches between CDSs of *A. svalbardicum* and of *A. pisum* were carried out with an e-value threshold of 10^-8^. The following steps were performed with an R script. Reciprocal best hit criterion was used to identify putative orthologous genes between the two species. These were aligned by the pairWiseAlignment function of the Biostrings package (Pages et al. 2016). Indels were inspected to flag CDS regions where the two species did not present the same reading frame. Bases in these regions were replaced with Ns, and were therefore trimmed by the Gblocks program (Castresana 2000; Talavera and Castresana 2007), alongside regions of unreliable alignment. We then estimated pairwise synonymous (*dS*) and nonsynonymous (*dN*) substitution rates for each gene, using the codon-based method of Li (1993), as implemented in the R package seqinR (Charif and Lobry 2007). Only the 9,696 genes (out of 9,924) with an alignment length of >90 nucleotides, *dN* <0.3 and *dS* <2 were kept. We also truncated *dN/dS* ratios to a maximal value of 2.5.

### Estimates of selection intensity based on intraspecific polymorphism

Polymorphism data for *A. pisum* was obtained from 60 genotypes originating from three *Medicago sativa* fields located in France and Switzerland (Jaquiéry et al. 2012b). These fields can be considered to harbour a single large population of pea aphids (Jaquiéry et al. 2012b, Peccoud et al. 2009a). DNA was individually extracted from four asexual females of each clone using the method described above. Because the approach described below does not require reconstructing allele sequences or individual genotypes, sequencing the pooled individuals (Gautier et al. 2013) was used to save costs. After RNAse treatment on each sample and DNA dosage with Pherastar, DNA samples were pooled to attain equimolar proportions. Paired-end libraries were then sequenced on two lanes of Illumina HiSeq 2000 using the Illumina Sequencing Kit v3 (producing 100-bp reads) by Beckman Coulter Genomics (Danvers MA, USA). This yielded ∼85X of sequencing coverage, hence an expectation of 0.71X per individual chromosome. Reads were mapped on Acyr 2.0 and symbiont genome sequences as described previously. The two alignment (BAM) files (one per sequencing lane) were filtered from PCR duplicates using SAMtools rmdup (Li et al. 2009) and reads realigned near indels using the Genome Analysis Toolkit (McKenna et al. 2010). The raw sequence data have been deposited in the SRA division of Genbank (project accession: PRJNA385905).

The two BAM files were merged and converted as pileup format using SAMtools (options -B -Q 0 –R) (Li et al. 2009). A modified estimator of Tajima’s D (Tajima 1989) which takes into account sequencing errors (Achaz 2008) was then calculated from this mpileup with Popoolation 1.2.2 (Kofler et al. 2011), after subsampling at a uniform coverage (subsample-pileup.pl, options: --target-coverage 30 --max-coverage 120 --method withoutreplace). Computations were performed for each gene including introns (Variance-at-position.pl --pool-size 120). Tajima’s D allows evaluating the type of selection at work, since selective sweeps and/or purifying selection tend to decrease it, and balancing selection tends to increase it.

The McDonald and Kreitman (1991) approach, which compares fixed mutations to polymorphic mutations in CDS, was adopted to further evaluate selection pressures on these different categories of genes, using the DoS estimate (Direction of Selection, Stoletzki and Eyre-Walker 2011). Positive, null and negative values of DoS respectively suggest adaptive evolution, neutral evolution, and purifying selection. Fixed mutations between species were counted from alignments we previously generated for *A. svalbardicum* and *A. pisum* CDSs. We restricted the analysis to regions of reliable alignments, as given by the Gblocks txts outputs. In these regions, we called SNPs on the BAM files with LoFreq (Wilm et al. 2012), which offers a good compromise between speed, sensitivity and accuracy in pools of multiple individuals (Huang et al. 2015). We used SAMtools mpileup (Li et al. 2009) to assess depth of coverage at all positions in these regions, polymorphic or not. We instructed mpileup to discard reads with mapping quality 0. The following was done in R. We discarded all positions covered by less than three reads (both BAM files combined). At each SNP, the number of polymorphic mutations was the number of different bases (alleles) found in the pea aphid population minus one. A fixed difference was counted if no base was shared at a position between the pea aphid population and *A. svalbardicum*. The number of polymorphic non-synonymous mutations per codon was taken as the number of amino acids found in the pea aphid populations for that codon minus one. To count the number of fixed non-synonymous differences per codon, we considered that a codon might differ between the two species by up to three mutations. Any of these may involve a change in protein sequence that we cannot ascertain without knowledge on the order of appearance of the mutations. We adopted parsimony and considered the minimum number of mutations required between the two codons. If several codons were present in the pea aphid population (due to a SNP), we considered the minimum number of coding changes that any pair of codons between the pea aphid and *A. svalbardicum* involves. For all these counts, we discarded rare codons showing more than one SNP, because the actual codons (and amino acids) present in the pea aphid population cannot be determined without phasing. We counted the following for each gene: the number of polymorphic non-synonymous changes (*Pn*), the number of all polymorphic changes minus *Pn* (which is the number of polymorphic synonymous mutations, noted *Ps*), the number of fixed non-synonymous differences (*Dn*), the number of all fixed differences minus *Dn* (which is the number of fixed synonymous changes, noted *Ds*). DoS was then calculated as *Pn*/(*Pn*+*Ps*) – *Dn*(*Dn*+*Ds*) by concatenating all genes from a given class of expression to avoid divisions by zero, due to genes without polymorphism or divergence, as done in Burgarella et al. (2015). We considered only genes whose average depth of coverage, as given by mpileup, was between 20 and 150 (expected coverage was ∼85) to avoid including genes presenting multiple collapsed copies that could artificially inflate polymorphism.

### Statistical analyses

Differences in expression levels between X-linked and autosomal genes in the different morphs, as well as differences in *dN/dS*, *dN* and *dS* between X-linked and autosomal genes were tested with Mann-Whitney U tests. The latter analysis was done on all genes, and on genes grouped based on average expression over the three morphs. To evaluate evolutionary forces responsible for faster-X evolution, we then compared *dN/dS*, Tajima’s D and DoS between X-linked and autosomal genes for classes of genes with different expression patterns (unbiased, male-biased, sexual female-biased and asexual female-biased genes). For biased genes, we considered different fold changes in expression (2-to 5-fold, and > 5-fold). Statistical significance was evaluated with Mann-Whitney U tests.

## Results

### Gene assignment to the X and autosomes

Based on the ratio of depth of coverage by reads from males over reads from females, 64% of the nucleotides assembled in the pea aphid reference genome (Acyr 2.0) were assigned to autosomes, 31% to the X chromosome while only 5% could not be assigned (supplementary figure S1). Genotypes of males at SNPs that were heterozygous in the female generally confirmed the assignment from coverage depth data (supplementary figure S2), though confirmation was not possible in regions lacking such SNPs. These estimates roughly correspond to the expected size of the X chromosome in the pea aphid, which represents ∼30% of the chromosome content based on karyotypes (Mandrioli and Borsatti 2007). This assignment revealed a very high rate of misassembly in Acyr 2.0: 56% of scaffolds of size ≥150 kb (which represent 80% of the assembly length) comprised blocks assigned to different chromosome types (supplementary figure S3 and table S1). Based on assigned scaffold blocks, 19,263 predicted genes were located on autosomes, 13,726 on the X chromosome while 4,001 genes could not be unambiguously assigned. The X chromosome contains a higher fraction of predicted genes than expected from its relative size (42%, test of proportion, *p*<10^-15^).

### Gene evolutionary rates

By comparing pea aphid gene sequences to transcripts sequenced from a related species (*A. svalbardicum*), we assessed the rates of substitution. Results confirm faster-X evolution, as X-linked genes present almost a twice higher non-synonymous substitution rate, *dN* (mean *dN*_X_= 0.034; *dN*_A_= 0.019, Mann-Whitney U = 4839860, *p* <10^-15^, *n* = 9096) and only slightly higher synonymous substitution rate, *dS* (mean *dS*_X_=0.101; *dS*_A_= 0.085, Mann-Whitney U = 6190137, *p* <10^-6^). As a result, the evolution of X-linked genes involves more protein-sequence changes (in proportion) than the evolution of autosomal genes (mean *dN*_X_*/dS*_X_ = 0.390; *dN*_A_*/dS*_A_ = 0.237; Mann-Whitney U = 5026170, *p* <10^-15^, figure 2A and supplementary table S2).

**Fig. 2.**
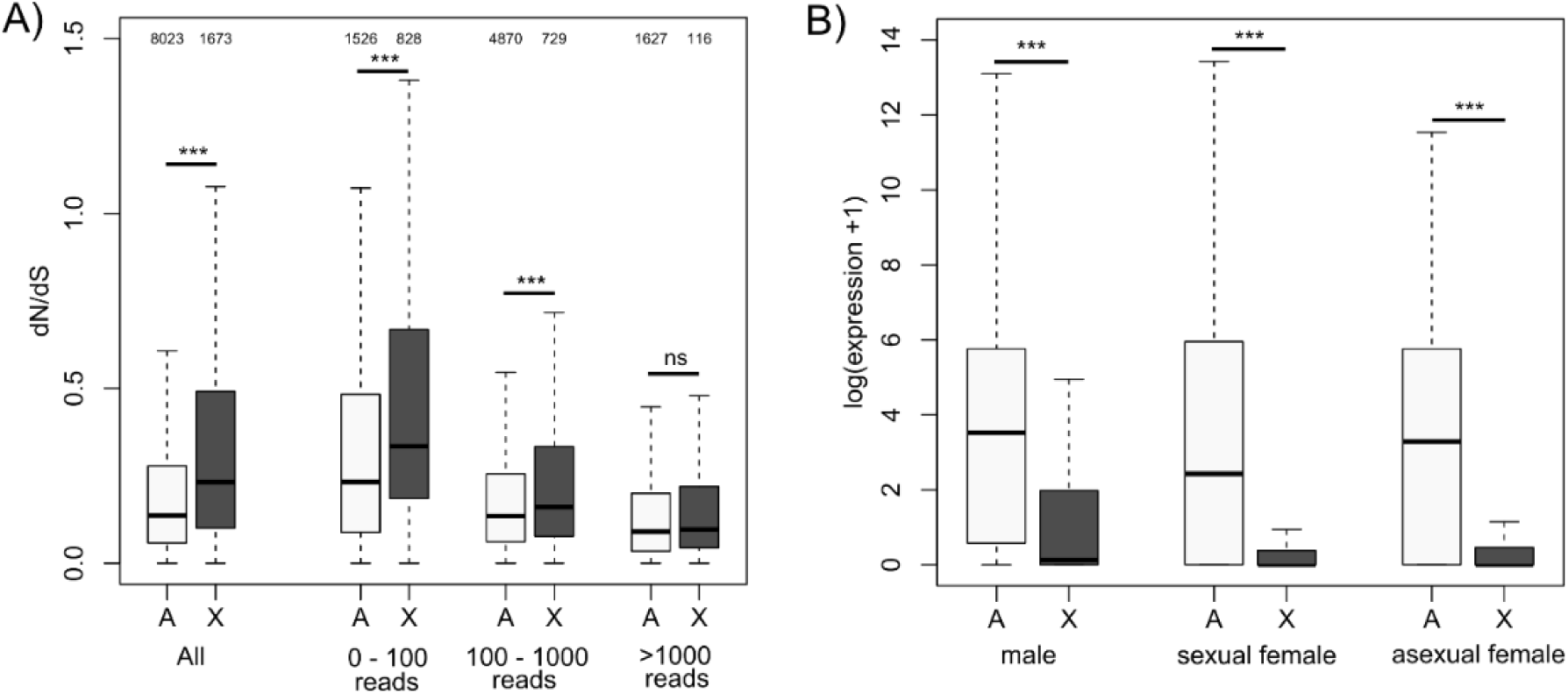
Evolutionary rates for autosomal and X-linked genes and gene expression in males, sexual and asexual females. A) Evolutionary rates (*dN/dS*) are shown for all genes (expressed or not in *A. pisum*) and for genes expressed at different levels (when averaged over male, sexual and asexual females): lowly expressed genes (i.e. covered by < 100 reads/kb of exon model); moderately expressed (from 100 to 1000 reads/kb), highly expressed genes (>1000 reads/kb). The number of genes per category is shown above each boxplot. B) Expression level for X-linked (*n* = 13613) and autosomal genes (*n* = 18812) in males, sexual females and asexual females. It should be noted that males carry only one X chromosome per cell and females carry two. Significance was tested with Mann-Whitney U tests.

### Causes of Faster-X evolution

High ratios of non-synonymous to synonymous rates, *dN/dS*, can result from a decreased influence of selection -and thus an increased influence of drift -on amino acid substitutions (i.e., relaxed negative selection) and/or from more efficient selection of adaptive changes in the protein sequence (i.e., increased positive selection). To balance these two hypotheses, we used the expression levels of genes as proxies for their possible impact on fitness. Expression levels of X-linked genes are lower than those of autosomal genes in all three main aphid morphs: males, sexual females and parthenogenetic females (figure 2B). Expression level averaged over the 13,726 X-linked genes ranges from ∼52 reads/kb in sexual and asexual females to 155 reads/kb in males, while expression level averaged over the 19,263 autosomal genes varies from 575 (in asexual females) to 648 reads/kb in males. Genes that are not expressed or expressed at a low level may have a reduced effect on the phenotype and may therefore accumulate non-synonymous substitutions faster (reduced purifying selection). We indeed observe that, for both X and autosomes, *dN/dS* ratios decrease with increasing expression levels (averaged over the three morphs), and that the contrast between X and autosomes tends to decline for highly expressed genes (figure 2A, supplementary table S2). Slight contrast in *dN* ratios is still maintained for highly expressed genes, though (supplementary table S2). Therefore, low expression levels of X-linked genes may not entirely account for faster X-evolution in aphids.

Selection may also be relaxed in genes that are predominantly expressed in rare morphs (males and sexual females), which constitute a minor fraction of the annual life cycle of aphids dominated by asexual females. Relaxed selection on mutations affecting male-biased genes (Purandare et al. 2014, Brisson and Nuzhdin 2008), combined with the tendency of such genes to locate on the X (Jaquiéry et al. 2013; Pal and Vicoso 2015) could contribute to faster-X evolution. However, the influence of X-linkage could not properly be evaluated in Purandare et al. (2014) because misassembled scaffolds, rather than individual genes, were assigned to chromosomes. Our new dataset of X-linked and autosomal genes unambiguously confirms that the X is largely enriched in genes over-expressed in males, and to a smaller extent in those over-expressed in sexual females (table 1). Like Purandare et al. (2014), we observe higher *dN/dS* ratios in genes over-expressed in the rarer morphs (i.e. males and sexual females, figure 3B-C), but not in genes over-expressed in the common morph (parthenogenetic females, figure 3D) when considering X-linked and autosomal genes together. Tajima’s D also tends to increase in genes over-expressed in the sexual morphs compared to unbiased genes (significantly so for all male-biased and for 2-to 5-fold female-biased genes, figure 4B), but not in genes over-expressed in the common morph (where D is significantly lower compared to unbiased genes, figure 4D), a pattern compatible with relaxed selection on genes expressed mainly in the rare morphs. However, the DoS did not differ significantly between these categories of genes, except for strongly female-biased genes (supplementary figure S4).

**Fig. 3.**
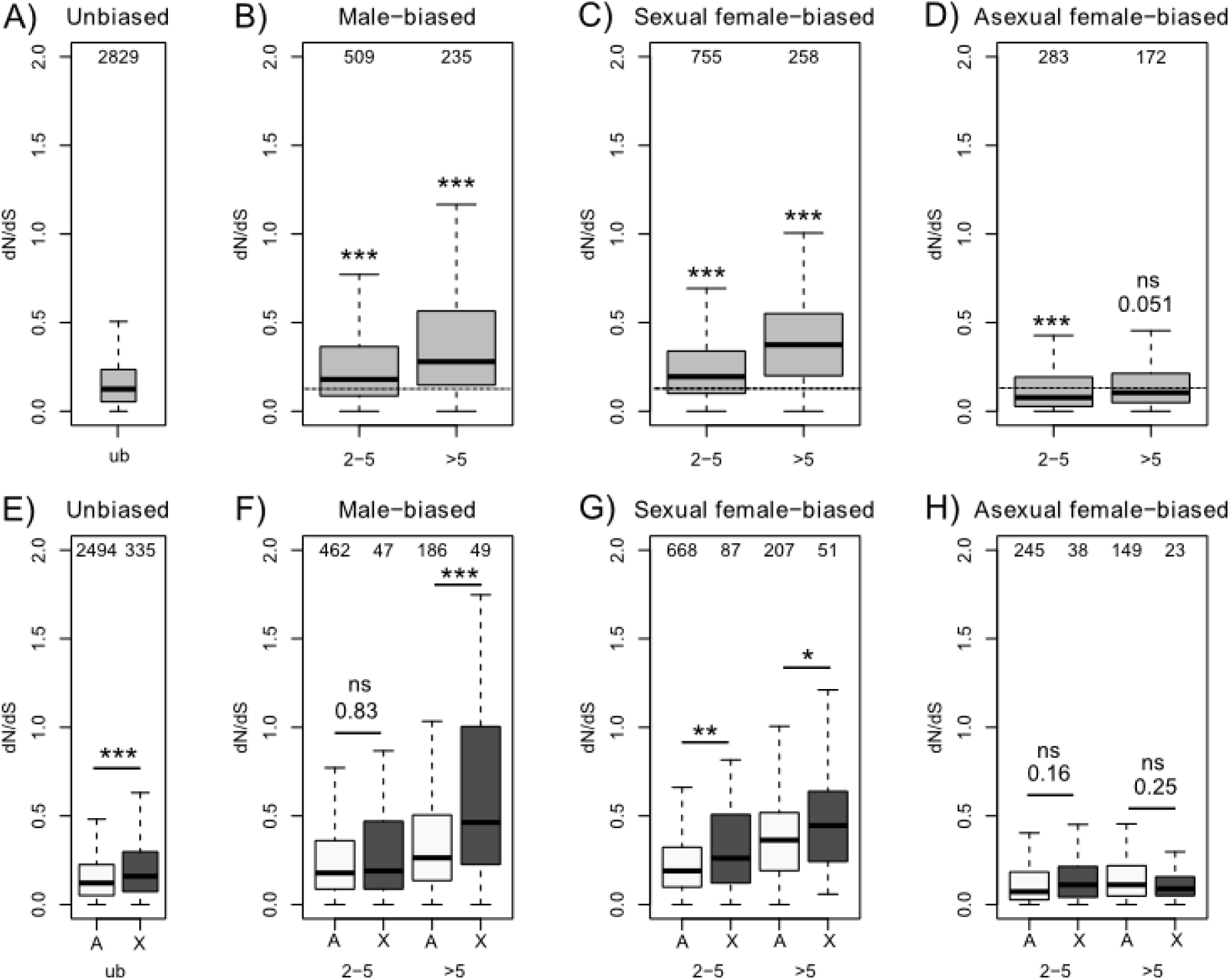
Substitution rates of genes (*dN/dS*, measured between *A.pisum* and *A. svalbardicum*) according to the ratios of expression levels between morphs. Panels A-D consider all genes together (X-linked and autosomal), and panels E-H consider X-linked (dark grey) and autosomal (light grey) genes separately. The number of genes in each class is shown above each boxplot. Only genes supported by at least 100 reads/kb of exon model were retained. ub: unbiased genes (padj > 0.1 for morph effect on expression), 2-5: levels of gene expression are two to five times higher in the specified morph (padj <0.05 for morph effect), >5: levels of gene expression are at least five times higher in the specified morph (padj <0.05). Significance of differences: ns: *p* > 0.05; \**p*< 0.05, \*\**p* < 0.01, \*\*\**p* < 0.001 (Mann-Whitney U tests). For panels B-D, differences correspond to comparisons with genes of the “unbiased” category, while X and autosomes were compared in panels F-H.

When analyses are done by chromosome type, *dN*/*dS* ratios of X-linked genes are significantly higher than those of autosomal genes for both sexual female and male-biased genes (figure 3E-3H), but not for asexual females. Contrastingly, Tajima’s D differs between chromosome types only for strongly male-biased genes (being lower for X-linked genes, suggesting more positive selection, figure 4F) and for unbiased genes (being larger for X-linked genes, possibly revealing more balanced selection, figure 4E). As a result, the enrichment of the X with genes expressed in the rare male morph (which have been hypothesized to evolve under relaxed selection) may not strongly contribute to faster-X evolution. No signal was detected between chromosome types based on the DoS index (supplementary figure S4).

**Fig. 4.**
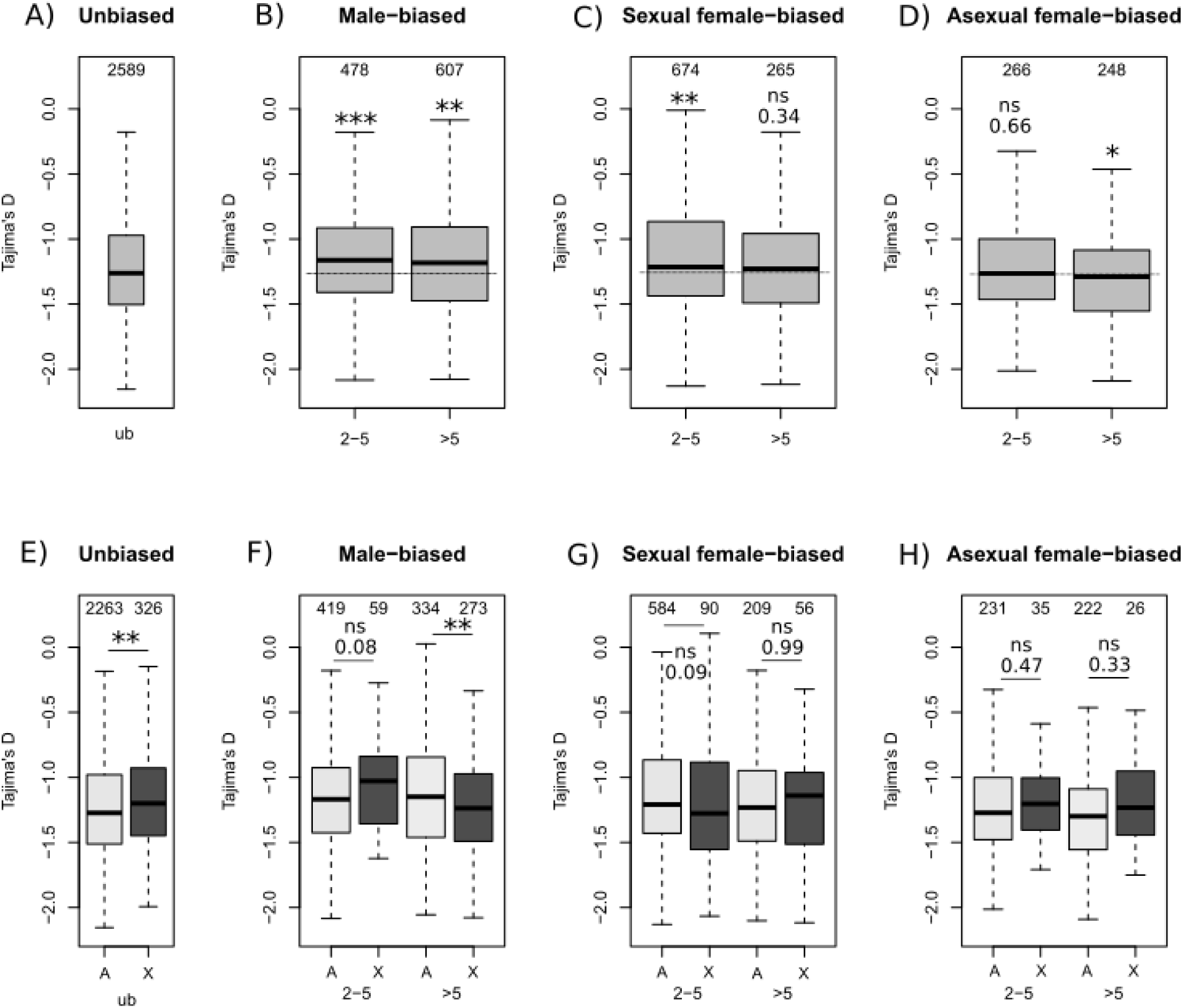
Tajima’s D according to the ratios of gene expression levels between morphs. Panels A-D consider all genes together (X-linked and autosomal) and panels E-H consider X-linked (dark grey) and autosomal (light grey) genes separately. Terms are defined as in Figure 3. Dashed lines show median values for unbiased genes. Significance of differences was tested with Mann-Whitney U tests.

Alternatively, the faster evolution of X-linked male-biased genes compared to autosomal male-biased genes could result from the fact that the former are present in hemizygous state in males. Non-synonymous mutations on the X are thus more exposed to selection in males, since they are not masked by other potentially dominant alleles, such that adaptive mutations on the X should more rapidly and more likely reach fixation than adaptive mutations on autosomes. This hypothesis predicts that the contrast between substitution rates of X-linked genes and autosomal genes will be highest for male-biased genes, and lowest for sexual-and asexual female-biased genes, because in these morphs, the X is always diploid and adaptive mutations can be recessive. We indeed observe these patterns (figure 3F), and the significantly lower Tajima’s D for X-linked male-biased genes compared to autosomal genes provides some support to this hypothesis (figure 4F).

## Discussion

Here we performed a genome-wide identification of X-linked genes enabling to locate a large number (13,726) and proportion (42%) of predicted genes on the X chromosome. We demonstrated that these genes tend to evolve faster than autosomal genes on average, confirming earlier results based on a much smaller set of coding genes (Jaquiéry et al. 2012a). Faster-X evolution mainly results from low expression of a large fraction of X-linked genes, which would less affect the phenotype and may accumulate non-synonymous mutations. Our analyses also suggest that higher exposure of recessive X-linked alleles to selection in hemizygous males might also partially contribute to faster-X evolution via more efficient positive selection.

We demonstrate clear faster-X evolution in the pea aphid based on a large set of X-linked and autosomal genes. The non-synonymous to synonymous substitution ratio (*dN/dS*) for X-linked genes is 1.69 times greater than for autosomal genes. This clearly places aphids among species showing strong contrast between evolution of X-linked and autosomal genes. Excluding sex-specific genes (e.g. Torgerson 2003; Kousathanas et al. 2014) and genes that escape post-meiotic silencing (Sin et al. 2012) for which X-linked sequences evolve much faster than autosomal ones, the *dN/dS* for X-linked genes is between ∼0.9 and ∼1.8 times that of autosomes in most species studied (i.e., *Drosophila*, mammals, birds and moths, review in Meisel and Connallon 2013; see also Sackton et al. 2014).

Remarkably, the pea aphid presents the same effective population size for the X and autosomes (Jaquiéry et al. 2012a), such that hemizygosity in males should be the only differentiating factor affecting the evolution of genes located on different chromosome types. However, our analyses revealed another key difference between X-linked and autosomal genes, in that the former are on average 4 to 10 times less expressed than the latter (figure 2B), and expectedly show higher rates of substitution. Such negative correlations between substitution rates and expression level have already been observed in several species (e.g., Drummond et al. 2005, Nguyen et al. 2015, Zhang and Yang 2015). Therefore, enrichment of the X with lowly expressed genes explains faster-X evolution in aphids to a large extent.

Gene expression differs between chromosome types in another dimension, as the X is enriched in genes that are mostly expressed in the rare morphs (i.e., males and sexual females) (Jaquiéry et al. 2013, Pal and Vicoso 2015). Such genes should also evolve under more relaxed constraints as they are exposed to the selective environment only during a short period of the aphid life cycle (Brisson and Nuzhdin 2008, Purandare et al. 2014). However, our analyses revealed that X-linked male-biased genes show lower Tajima’s D than autosomal male-biased genes (figure 4F-4G). As a result, enrichment of the X chromosome with genes expressed in the rare male morph is unlikely to be an important contributor to faster-X evolution in the pea aphid.

This leaves hemizygosity of the X chromosome in males, which exposes all X-linked alleles expressed in males to the selective environment, as another contributing cause. This hypothesis is supported by the contrast in *dN/dS* between X-linked genes and autosomal genes, which is larger for male-biased genes than for sexual and asexual females-biased genes (figure 3F-3H), and by the lower Tajima’s D on male-biased X-linked genes than for those on autosomes (figure 4F-4G), but not by the DoS, which shows no sign of selection.

Mean expression levels of X-linked genes measured in the whole body are strikingly lower than those of autosomal genes, in all morphs studied. The difference we found is more pronounced than in previous observations (Jaquiéry et al. 2013, Pal and Vicoso 2015) probably due to more reliable gene assignments to chromosomes (we found that 10% of the 3,712 genes used in Jaquiéry et al. 2013 had been misassigned because of scaffold misassembly). Lower expression of X-linked genes compared to autosomal genes is observed in mammals (Nguyen et al. 2015), but not in *Drosophila* (Zhang and Presgraves 2016). To our knowledge, no other taxon displays such a strong contrast between the X and autosomes gene expression levels as does the pea aphid. This raises the question as to why genes on the X are so little expressed in this species. A theoretical model (Jaquiéry et al. 2013) predicts that the X chromosome is more easily invaded by sexually antagonistic alleles beneficial to males and deleterious to females than autosomes. This may have favored a global decrease in gene expression of this chromosome in the common morph (the asexual females) for which it could be harmful. Pseudogenisation on the X chromosome would have ensued if genetic variation in lowly expressed genes has little effect on fitness. Yet, this chromosome carries one third of the genome and contains a higher fraction of genes than predicted by its relative size. Insights into the role of sexual antagonism on the peculiar expression patterns observed here could be gained by studying gene expression of this chromosome in different male and female tissues. Particularly, transcriptomes of tissues subject to contrasted sex-specific selection pressures (e.g. Parisi 2003; Khil et al. 2004; Yang et al. 2006; Huylmans and Parsch 2015) could help examine this hypothesis.

Assignments of scaffold blocks to chromosomes revealed widespread errors in the pea aphid genome assembly (Acyr 2.0). More than half of scaffolds larger than 150kb are clear chimeras of X and autosomes. This is a minimal estimate for the rate of misassembly, since our method only detects breakpoints between X and autosomes. Given that the X represents 30% of the pea aphid genome, assembly errors involving two X-linked genomic regions, and those involving two autosomal regions should respectively represent 0.3^2^ and 0.7^2^ (58% in total) of all assembly errors, a significant number of events that we could not detect.

Moreover, assembly errors involving fragments of less than 10 kb could not be detected given the resolution of our analyses (see methods). Other lines of evidence argue for frequent errors in the current pea aphid genome assembly. First, a genetic map of 305 microsatellite markers (Jaquiéry et al. 2014) validates our assignment to chromosomes, and also demonstrates high rates of scaffold misassembly (see supplementary file S1). Second, X-A breakpoints co-localize with sharp variations in genetic differentiation (*F*_ST_) between pea aphid populations from different host plants, as determined by high-density genome scans (Nouhaud et al. submitted). Third, synteny between Acyr 2.0 scaffolds and scaffolds from two *Myzus persicae* genomes (Mathers et al. 2017, available at www.aphidbase.com) stops at X-A breakpoints (data not shown). Fourth, entire scaffolds assigned by Bickel et al. (2013) to the X chromosome account for 11% of Acyr 2.0, much lower than the ∼30% of the genome that this chromosome represents, based on karyotypes (Mandrioli and Borsatti 2007). Bickel et al. (2013) explained this discrepancy by assembly errors combined with methodological bias towards assigning misassembled scaffolds to autosomes (a hypothesis we confirm by the comparison of scaffold assignments to chromosomes by Bickel et al. 2013 and the present study, see supplementary figure S5). By contrast, our new assignments to chromosomes provide estimates corresponding to the expected relative size of the X. Consequently, we confidently conclude that the genome of the pea aphid presents considerable assembly problems, to a degree that goes far beyond what current assembly pipelines typically yield (e.g. Salzberg et al. 2004; Muggli et al. 2015). While the cause of these errors remains undetermined, they have important drawbacks for genomic studies on a species that is currently considered the model aphid, in particular those relying on the physical organization of the genome, ranging from high-resolution genome scans to studies of chromatin conformation, genomic rearrangements, etc. We therefore urge for a reassembly of the pea aphid genome based on additional data. The results presented here should however not be affected by misassembly as we were able to assign unambiguously almost 90% of the 36,990 predicted gene of the pea aphid. This means that at fine-grain level, that of a few kilobases to few tens of kilobases (the range within which most genes are contained), the chimerism problem is not a major issue.

In conclusion, we have shown here that faster-X evolution of proteins in the pea aphid is principally explained by relaxed selection on lowly expressed genes, a class of genes that is more frequent on the X chromosome than on autosomes. Exposure of X-linked recessive alleles to selection in hemizygous males could play an additional, yet marginal, role in faster-X evolution. Importantly, the pea aphid complex, which includes races and cryptic species at different stages of divergence (Peccoud et al. 2009a, 2009b, 2014), offers excellent opportunities to investigate mechanisms driving genome evolution through comparative genomics. In particular, characterizing gene expression in morphs of closely and distantly related species could help disentangling the role of drift and selection in the low expression of the X and its masculinization.

## Acknowledgements

This work was supported by the Agence Nationale de la Recherche (grants ANR-09-GENM-017-001 to Denis Tagu, ANR-11-BSV7-005-01 to Denis Tagu and ANR-11-BSV7-007 to JCS); INRA-AIP BioRessources (project Poly-Express to JCS and Denis Tagu); the Fondation pour la Recherche sur la Biodiversité (grant AAP-IN-2009-020 to JCS); the Institut Polaire Francais Paul-Emile Victor (IPEV project 426 Arctaphid to JCS); the Genoscope (project 62 AAP 2009/2010 to JCS) and the Swiss National Science Foundation (grants PBLAA-122658 & PA00P3-139720 to J.J.). We would like to thank the GenOuest platform for the use of its cluster to perform the computational analyses.

